# Elexacaftor/VX-445-mediated CFTR interactome remodeling reveals differential correction driven by mutation-specific translational dynamics

**DOI:** 10.1101/2023.02.04.527134

**Authors:** Minsoo Kim, Eli Fritz McDonald, Carleen Mae P. Sabusap, Bibek Timalsina, Disha Joshi, Jeong S. Hong, Andras Rab, Eric J. Sorscher, Lars Plate

## Abstract

Cystic fibrosis (CF) is one of the most prevalent lethal genetic diseases with over 2000 identified mutations in the cystic fibrosis transmembrane conductance regulator (CFTR) gene. Pharmacological chaperones such as Lumacaftor (VX-809), Tezacaftor (VX-661) and Elexacaftor (VX-445) treat mutation-induced defects by stabilizing CFTR and are called correctors. These correctors improve proper folding and thus facilitate processing and trafficking to increase the amount of functional CFTR on the cell surface. Yet, CFTR variants display differential responses to each corrector. Here, we report variants P67L and L206W respond similarly to VX-809 but divergently to VX-445 with P67L exhibiting little rescue when treated with VX-445. We investigate the underlying cellular mechanisms of how CFTR biogenesis is altered by correctors in these variants. Affinity purification-mass spectrometry (AP-MS) multiplexed with isobaric Tandem Mass Tags (TMT) was used to quantify CFTR protein-protein interaction changes between variants P67L and L206W. VX-445 facilitates unique proteostasis factor interactions especially in translation, folding, and degradation pathways in a CFTR variant-dependent manner. A number of these interacting proteins knocked down by siRNA, such as ribosomal subunit proteins, moderately rescued fully glycosylated P67L. Importantly, these knock-downs sensitize P67L to VX-445 and further enhance the correction of this variant. Our results provide a better understanding of VX-445 biological mechanism of action and reveal cellular targets that may sensitize unresponsive CFTR variants to known and available correctors.

## Introduction

Cystic fibrosis (CF) is a common genetic disorder that affects about 32,000 people in the United States and 100,000 worldwide^1,2^. CF occurs due to mutations in the cystic fibrosis transmembrane conductance regulator (CFTR) protein^3^, a cyclic adenosine monophosphate (cAMP) dependent anion channel that conducts chloride and bicarbonate across epithelial apical membranes of multiple exocrine organs. In the respiratory tract, loss of CFTR results in lack or dysfunction of mucociliary transport leading to thick mucus buildup in lungs, reduced airflow, respiratory obstruction, and chronic infection^4,5^. Similarly, patients experience obstruction and malabsorption in the digestive system frequently with pancreatic insufficiency resulting in malnutrition and CF-associated diabetes^6^. Until now, over 2000 identified mutations and polymorphisms have been identified in the CFTR gene but research for therapeutic interventions has been primarily centered on the F508del variant presented in approximately 90% of patients^7^. Novel corrector drugs, such as Elexacaftor (VX-445) in combination with other CFTR modulators, are highly effective in individuals with F508del but demonstrate a variable response in rarer variants which has impeded FDA approval for a number of molecular defects^8,9^. Understanding the precise mechanism that underlies correction of a particular variant is crucial for development of personalized CF therapy.

Preceding VX-445, pharmacological correctors such as Lumacaftor (VX-809) and Tezacaftor (VX-661) were approved for use in combination with the CFTR ion transport potentiator Ivacaftor (VX-770) due to efficacy in correcting the F508del mutation^10–13^. Correctors help rescue many class II CFTR mutations that cause impaired protein folding and targeted degradation by endoplasmic reticulum associated degradation (ERAD) pathway^14^. Hence, correctors partially restore defective protein trafficking and increase steady-state CFTR levels at the plasma membrane. VX-809 and VX-661 are type I correctors that stabilize the protein interface between nucleotide binding domain 1 (NBD1) and transmembrane domain 1 (TMD1)^15–17^, and share a common binding site in TMD1^18^. By contrast, VX-445 represents a distinct pharmacophore with more complex structural features compared to VX-809 or its analog VX-661. VX-445 binds at the interface of transmembrane helices 2, 10, 11 and the lasso motif^19^ to allosterically stabilize NBD1 termed a type III corrector^20^. These type I and type III correctors, when applied in combination, synergistically correct CFTR folding defects likely due to distinct mechanisms of action^21^. Despite their efficacy in the clinic, current corrector therapies are reported to cause side effects including diarrhea, chest tightness, neurocognitive adverse events and interference with other drugs^9,22,23^. In addition, patients with rare mutations that have not yet been approved for application of VX-445 are left with less-effective conventional CF treatments. These complications underline the importance to mechanistically understand CFTR mutation specific modulator responses.

Potential factors implicated during CFTR rescue by correctors have included cellular components and pathways such as ER quality control, cytoskeleton organization and immune response^24–26^. However, CF phenotype likely manifests from additive derailments of protein homeostasis (proteostasis) pathways. In this context, CFTR biogenesis is highly regulated at multiple stages by proteostasis factors that govern translation, translocation, molecular chaperones and other folding machinery^27–30^. Once misfolded, degradation pathways direct ubiquitinated CFTR to the proteasome. Properly folded CFTR undergoes several quality-control checkpoints to be trafficked to the Golgi and the plasma membrane where it is recycled or degraded through endocytosis and lysosomal metabolism. In a profiling study of the F508del CFTR proteostasis network, F508del showed perturbations across numerous proteostasis factors in distinct pathways compared to wild-type (WT) CFTR, indicating a mutation can impact multiple protein-protein interactions^31^. Similarly, correctors such as VX-809 that counteract mutation-induced structural defects can revert aberrant protein interactions of F508del and restore CFTR protein interactions to more closely reflect those of WT^32,33^. These studies provide evidence that mass spectrometry proteomics analysis is suitable for systematic and sensitive measurement of alterations in the CFTR proteostasis network and provides information regarding the basis of biological mechanisms that underlie CFTR pharmacological correctors.

Here, we applied tandem mass tag (TMT)-multiplexed affinity purification-mass spectrometry (AP-MS) to elucidate ways in which CFTR corrector treatment is influenced by mutation-specific interaction partners. We found the CF causing P67L variant to be a VX-809 selective responder which was minimally rescued by VX-445, whereas the L206W variant is an additive responder (rescued by either VX-809 or VX-445). We then measured proteostasis changes of P67L and L206W in response to the two clinically validated modulators. VX-809 partially reverts the interactome of P67L and L206W towards that of WT, whereas VX-445 fails to restore early-stage proteostasis interactions with regard to translation, folding, or degradation machineries in the P67L variant. These findings indicate that VX-445 acts after CFTR passes early quality control checkpoints such as translation and folding. We also demonstrate that siRNA knock-down of select ribosomal proteins results in the enhanced rescue of mature P67L by VX-445, indicating that unresponsive class-II mutants can be sensitized to VX-445 by modulating translational pathways.

## Results

### I. Rare CFTR mutants P67L and L206W demonstrate distinct corrector responses

Class II mutations impair conformational maturation of CFTR and lead to degradation by ERAD^34^. Unlike F508del, CFTR variants such as P67L and L206W are markedly responsive to VX-809-leading to their designation as *hyper-responsive*^33,35^. We investigated these two hyper-responsive mutations by transiently transfecting HEK293T cells with mutant CFTR constructs and treating cells with CFTR correctors VX-809, VX-661, and VX-445 (**Figure 1A**). P67L showed robust correction by VX-809 or VX-661 as measured by C-band expression indicative of folded and fully glycosylated CFTR at the plasma membrane. In contrast, VX-445 exhibited nominal rescue of P67L, showing no significant difference from control (DMSO). Treatment with both VX-445 and VX-661 led to a response similar to VX-661 alone, further corroborating the minimal correction effect of VX-445 on P67L. On the other hand, all individual correctors strongly rescued trafficking of L206W, despite this CFTR variant being an early TMD1 mutation like P67L (**Figure 1B and 1C**). Differential response to VX-445 by these two hyper-responsive mutations suggests important differences in proteostasis that govern CFTR correction.

**Figure 1.**
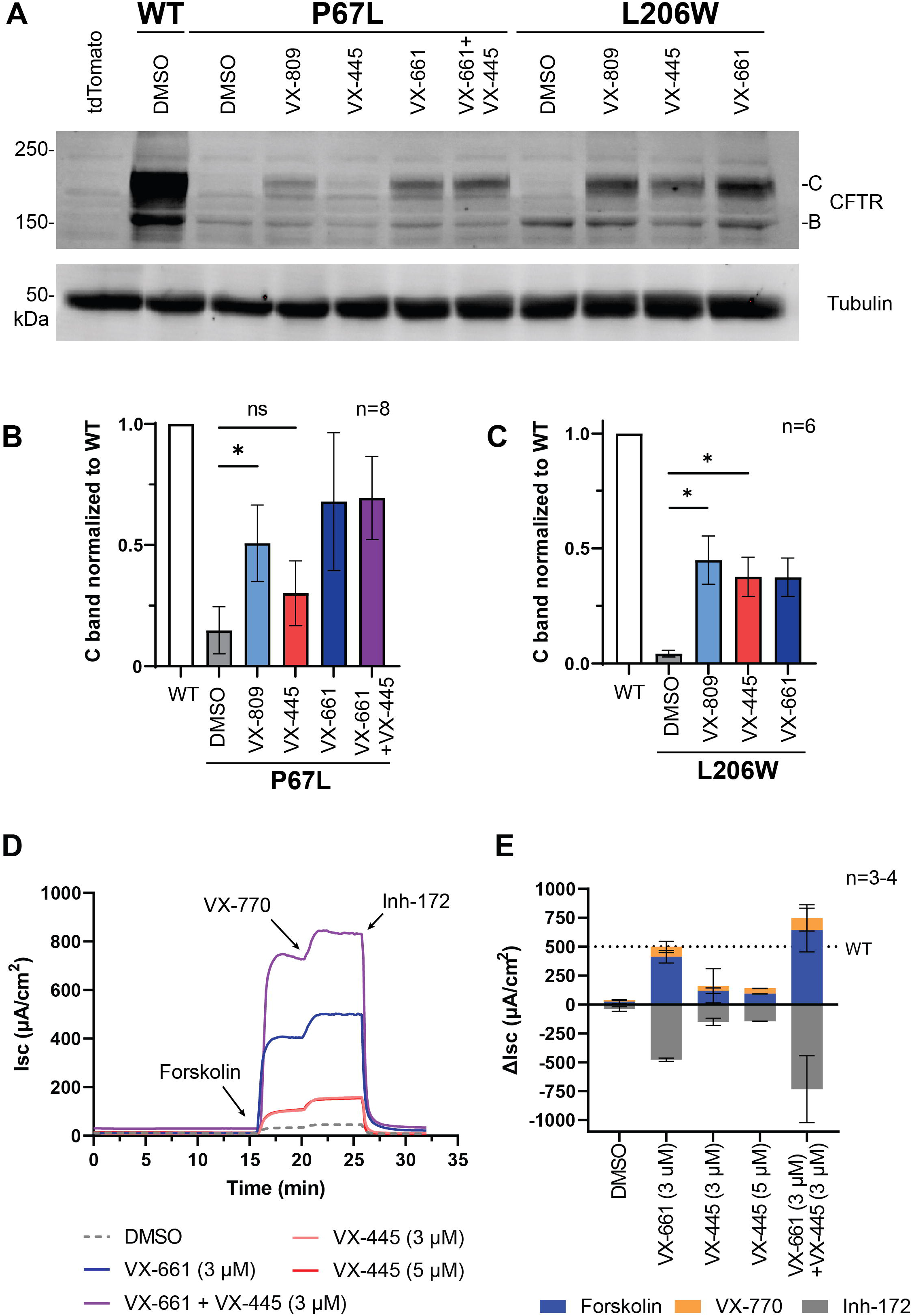
P67L CFTR is not responsive to VX-445. **A.** Representative Western blot image showing expression of CFTR in HEK293T cells transiently transfected with tdTomato control or CFTR contracts (WT, P67L, and L206W) in response to drug treatments (3 μM) for 24 h. P67L CFTR is restored to near wild-type levels with VX-809 treatment whereas VX-445 treatment remains indistinguishable from control. L206W is rescued by all three correctors. **B.** Quantification of P67L CFTR C bands from A shown as mean ± SEM. Statistical differences were computed via oneway ANOVA with Geisser-Greenhouse correction and post-hoc Tukey multiple comparisons testing within each mutant (p-values: * < 0.05, ** < 0.01, *** < 0.001, and **** < 0.0001). **C.** Quantification of L206W CFTR C bands from A shown as mean ± SEM. Statistical analysis as in B. **D.** Ussing chamber measurements on FRT cells stably expressing P67L. Data showing short-circuit current (Isc) after luminal stimulation with forskolin (5 μM) and VX-770 (5 μM) and inhibition by CFTR inh-172 (10 μM) While VX-661 restores ion current close to WT level, VX-445 restoration remains at low levels at both concentrations (3 μM and 5 μM). **E.** Bar graph representation of the acute treatment induced Isc changes expressed as mean ± SD across 3-4 biological replicates.

To confirm that the biochemical findings translate to functional output, we performed Ussing chamber analysis on the two variants after interventions with diverse correctors. Fischer Rat Thyroid (FRT) cells stably expressing P67L or L206W CFTR were treated individually with VX-661 or VX-445, or in combination. Briefly, cells were exposed sequentially to forskolin to induce channel opening, followed by CFTR potentiator VX-770 and inhibitor-172 to test effects on short circuit current (Isc). Similar to Isc current improvement of P67L following VX-809 treatment shown previously^36^, VX-661 restored channel function close to WT level (**Figure 1D and1E**). VX-445 mildly corrected the function of P67L variant compared to vehicle treatment only reaching 30% of VX-661 correction. Two concentrations of VX-445 at 3 μM and 5 μM showed that the correction by this compound is saturated, unlike what has been shown with VX-661 and VX-809 previously^37^. This further supports P67L as a selective responder to VX-809 while remaining only mildly responsive to VX-445. In contrast, function of L206W CFTR was equally restored by VX-661 or VX-445 comparable to WT level in a dose dependent manner (**Figure S1**).

Having demonstrated the distinct responsive properties of P67L and L206W by both biochemical and electrophysiology assays, we next aimed to define the proteostasis network for these mutants in response to corrector drugs, including distinct proteostasis pathways that which may drive the differential response to VX-445.

### II. VX-445 does not redirect P67L protein-protein interactions towards WT profile

To quantitatively investigate the underlying proteostasis factors driving differential corrector rescue, we performed LC-MS/MS coupled with isobaric tandem mass tag (TMT) labeling on co-immunoprecipitated CFTR and its interactors (**Figure 2A**). This workflow was based on previous studies that elucidated the proteostasis interactome changes of mutant thyroglobulin^38^ and mutant CFTR in response to correctors^33^. Briefly, HEK293T cells were transfected with WT, P67L, or L206W CFTR constructs and treated with correctors VX-809 and/or VX-445. Cells were lysed and CFTR interacting proteins were co-immunoprecipitated using CFTR antibody (clone 24-1) conjugated beads. Proteins were digested to peptides that were then labeled with 11-plex TMT labels to multiplex all conditions into a single MS run. From the resulting tandem MS spectra of six individual biological replicates (**Table S1**), TMT reporter fragments were deconvoluted to compare intensities directly and quantitate the corresponding protein abundance across conditions.

**Figure 2.**
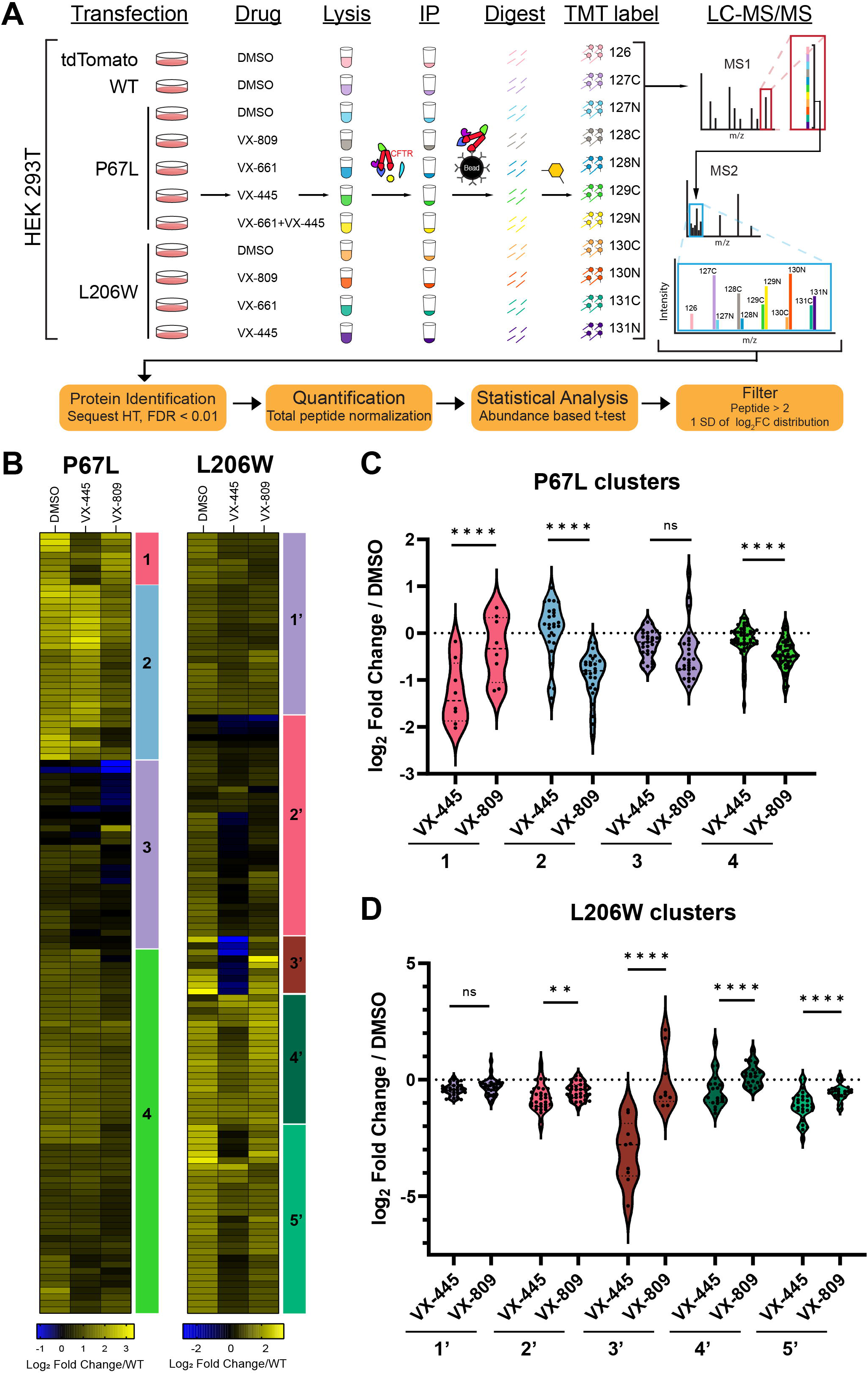
CFTR proteostasis interactome profiling via quantitative mass spectrometry analysis reveals attenuated protein interactions in effective drug treatment conditions. **A.** Multiplexed affinity purification-mass spectrometry workflow. Cells are transfected, drug treated, and lysed for coimmunoprecipitation (IP). Eluted proteins are digested with trypsin and TMT-labeled for pooled LC-MS/MS analysis to identify and quantify peptides. **B.** Hierarchical clustering of interactors grouped based on intensity similarity where WT CFTR normalized abundances were converted to an Euclidean distance matrix and clustered using Ward’s minimum variance method. Each bar is an interacting protein and color indicates log_2_ fold change in interaction with CFTR when compared to WT. For P67L, blue [2] and green [4] clusters represent interactions decreased by VX-809 whereas pink and red clusters [1, 2’ 3’] represent interactions decreased by VX-445 showing corrector-driven unique interactome changes. **C.** Violin plots of clusters in P67L CFTR show comparison of log_2_ fold changes of corrector treatment over DMSO. Paired two -tailed t-tests were performed within each cluster between VX-445 and VX-809 (p-values: * < 0.05, ** < 0.01, *** < 0.001, and **** < 0.0001). **D.** Violin plots of clusters in L206W CFTR show comparison of log_2_ fold changes of corrector treatment over DMSO. Statistical analysis as in C.

First, we prioritized a list of identified CFTR interactors for quantitative comparison. Interacting protein abundances were compared against a control mock transfection to determine confident CFTR interactors (**Figure 2A and Figure S2**). We filtered for interactors displaying a log_2_ fold enrichment (CFTR/control) greater than one standard deviation based on the distribution of identified proteins. Each CFTR variant and corrector treatment was compared to the control and a master list of confident CFTR interactors for further analysis (**Table S2**). Next, we normalized interactor abundances to WT CFTR and compared each mutant with and without drug treatment against WT, followed by hierarchical clustering was performed of the WT-normalized abundances to observe treatment-specific aberrant interaction behaviors (**Figure 2B**).

As described previously^33^, both mutants overall showed increased interactions that are potentially triggered by misfolding of the protein and enhanced recruitment of proteostasis factors. These aberrant interactions decreased in abundance with VX-809 treatment, reverting towards the WT interactome profile. These changes are reflective of restored folding of mutant CFTR, a pattern also observed previously^33^. VX-445 treatment for P67L failed to revert many interactions back to WT and more closely resembled control treatment (DMSO) (**Figure 2B and 2C** blue [2] and green [4] clusters). In contrast for L206W, VX-445 treatment mostly restored interactions close to or below WT levels. (**Figure 2B and 2D**). These distinct interactomics profiles are in line with the differential trafficking and functional results.

Also identified were groups of interactions uniquely impacted by VX-445 in both mutants indicating that each corrector can indeed influence discrete components of the proteostasis network (**Figure 2B-D** pink and red clusters [1, 2’, 3’]). Furthermore, we evaluated the combination treatment of VX-661 and VX-445, which was effective for correcting P67L. This combination showed restoration of interactions closer toward WT, consistent with defects in P67L not corrected by VX-445 alone that are impacting the proteostasis profile (**Figure S3).**

Overall, proteostatic, trafficking, and electrophysiology evidence suggest a divergent response of P67L and L206W to the type III corrector VX-445, despite both variants responding similarly to VX-809/661. We therefore sought to use our quantitative proteomics measurements to reveal the underlying proteostasis states and pathways driving this differential drug response.

### III. VX-445 treatment leads to distinct proteostasis states for P67L and L206W

To understand which proteostasis pathways differentially impact P67L and L206W drug response, interactors were classified into the following cellular pathways relevant to CFTR biogenesis: translation, folding/degradation, trafficking, cytoskeleton, and endocytosis (**Figure 3A**). Early biogenesis pathway interactions with P67L such as translation and folding/degradation failed to change with VX-445 treatment compared to control (**Figure 3B**). By contrast, these early pathway interactions showed significant decrease with VX-809, VX-661, and VX-445 and VX-661 combination treatments (**Figure 3B**). In L206W, VX-445 significantly decreased interactions in these early pathways (**Figure 3C**). The results therefore suggest mechanisms as early as translation, folding and degradative quality control are key checkpoints for pharmacological correction by VX-445.

**Figure 3.**
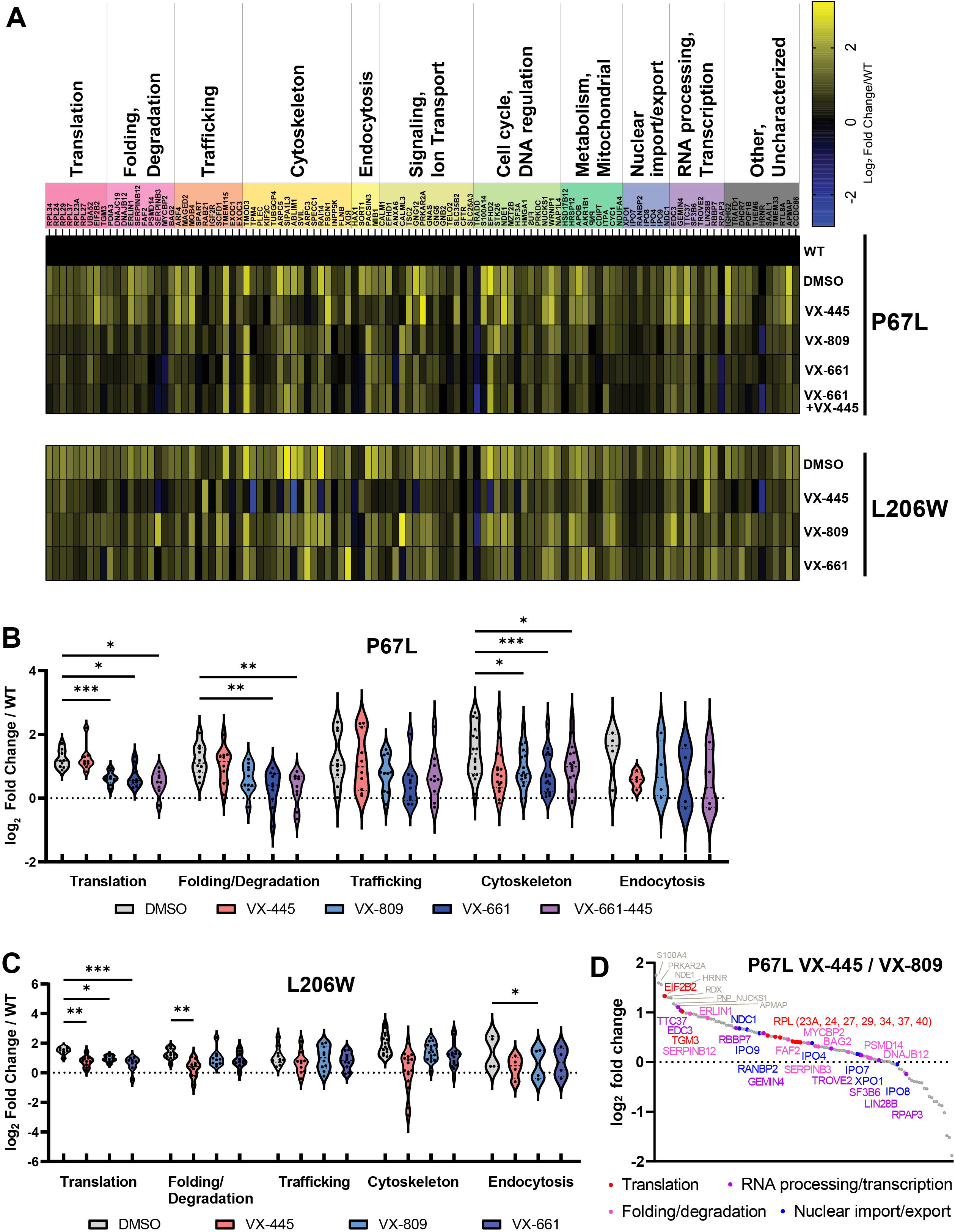
Pathway specific disruptions with translation and degradation in P67L. **A.** Translation and degradation factors show increased interactions with P67L that are restored toward WT with VX-809 but not VX-445. Heatmap of interactors classified into corresponding pathway categories. **B.** Violin plot of P67L interactors involved in CFTR processing pathways. Abundance of interactions of proteins in the two pathways were statistically unchanged in P67L treated with VX-445 when compared to control. One-way ANOVA with Geisser-Greenhouse correction and post-hoc Tukey multiple comparisons testing was performed within each pathway group (p-values: * < 0.05, ** < 0.01, and *** < 0.001). **C.** Violin plot of L206W interactors involved in CFTR processing pathways. Abundance of interactions of proteins were generally decreased by all treatments compared to control. Statistical analysis as in B. **D.** Waterfall plot directly comparing the interactions changes between VX-445 to VX-809 treatment for P67L. Higher log2 fold change indicates higher interaction in VX-445 treated P67L than that of VX-809 treated P67L. Shown in red are interactors involved in translation, and in purple are interactors involved in folding and degradation. Interactors that were over log2 fold change of 1 but did not belong in the highlighted pathways are noted in grey. Candidates such as EIF2B2 and RPLs were selected for follow up siRNA KD studies for further validation.

Next, we directly compared interactions that may govern distinct drug responses by examining a waterfall plot of log_2_ fold changes comparing VX-445 over VX-809 treatment (**Figure 3D**). As expected, interaction intensities of several proteins from the two disrupted pathways (translation, folding/degradation) such as eIF2B2, SERPINB12, ERLIN1, and several large ribosomal subunit proteins (RPLs: 23A, 24, 27, 29, 34, 37, 40) showed high log_2_ fold changes (**Figure 3D**). In contrast, we observed these proteins to be randomly dispersed in the L206W waterfall plot comparing VX-445 over VX-809, indicating their association with the distinct P67L drug response (**Figure S4**). We next sought to evaluate the relevance of these interactions for CFTR trafficking via siRNA knock-down and explore their impact on drug response.

### IV. Corrector mechanism is distinct for proteostasis interactions

We hypothesized that the measured proteostatic differences between P67L and L206W may be important for the pharmacological correction in these variants. To interrogate the role of proteostasis factors on CFTR trafficking, siRNA knock-down (KD) was performed in a Cystic Fibrosis Bronchial Epithelial cell-line with inducible P67L CFTR expression (TetON P67L CFBE)^39^. First, we selected a panel of targets in the pathways including translation, degradation, nuclear import/export, and RNA processing, from the waterfall plot (**Figure 4A and 4D**). The full list of the prioritized siRNA panel is reported along with siRNA KD confirmation with quantitative reverse transcription PCR (RT-qPCR) (**Table S3 and Figure S5**). Interestingly, siRNA KD of translation factor targets such as RPL34 and RPL37 mildly increased the amount of P67L band C under basal conditions indicating that these targets may ultimately promote degradation/sequestering of misfolded P67L CFTR (**Figure 4A-B**). We also treated cells with VX-445 after siRNA KD to investigate the effect of these KDs on drug response. Strikingly, we observed significant sensitization of P67L CFTR to VX-445 in several RPL KDs such as RPL24, 34, and 37 where mature CFTR (C band) was rescued close to levels observed with VX-809 (**Figure 4C**). Trafficking efficiency, as measured by calculating the ratios of band C/(C+B), was also improved with VX-445 after KD of RPL23A, 24, 29, 34, and 37 (**Figure S6**). Subsequently, factors involved in degradation, RNA processing and transcription were probed by siRNA KD (**Figure 4D**). Depletion of RBBP7 moderately rescued P67L band C (**Figure 4E**), while KD of serine-protease inhibitor SERPINB12 and RNA-binding protein RBBP7 induced significant sensitization to VX-445 (**Figure 4F**). These data highlight that translational protein interactions hinder correction by VX-445 either by directly interfering with binding or negatively impacting specific translational dynamics required for VX-445 drug response. Moreover, serine-protease inhibitors and epigenetic regulation in CFTR may hold previously unknown impact on pharmacological correction.

**Figure 4.**
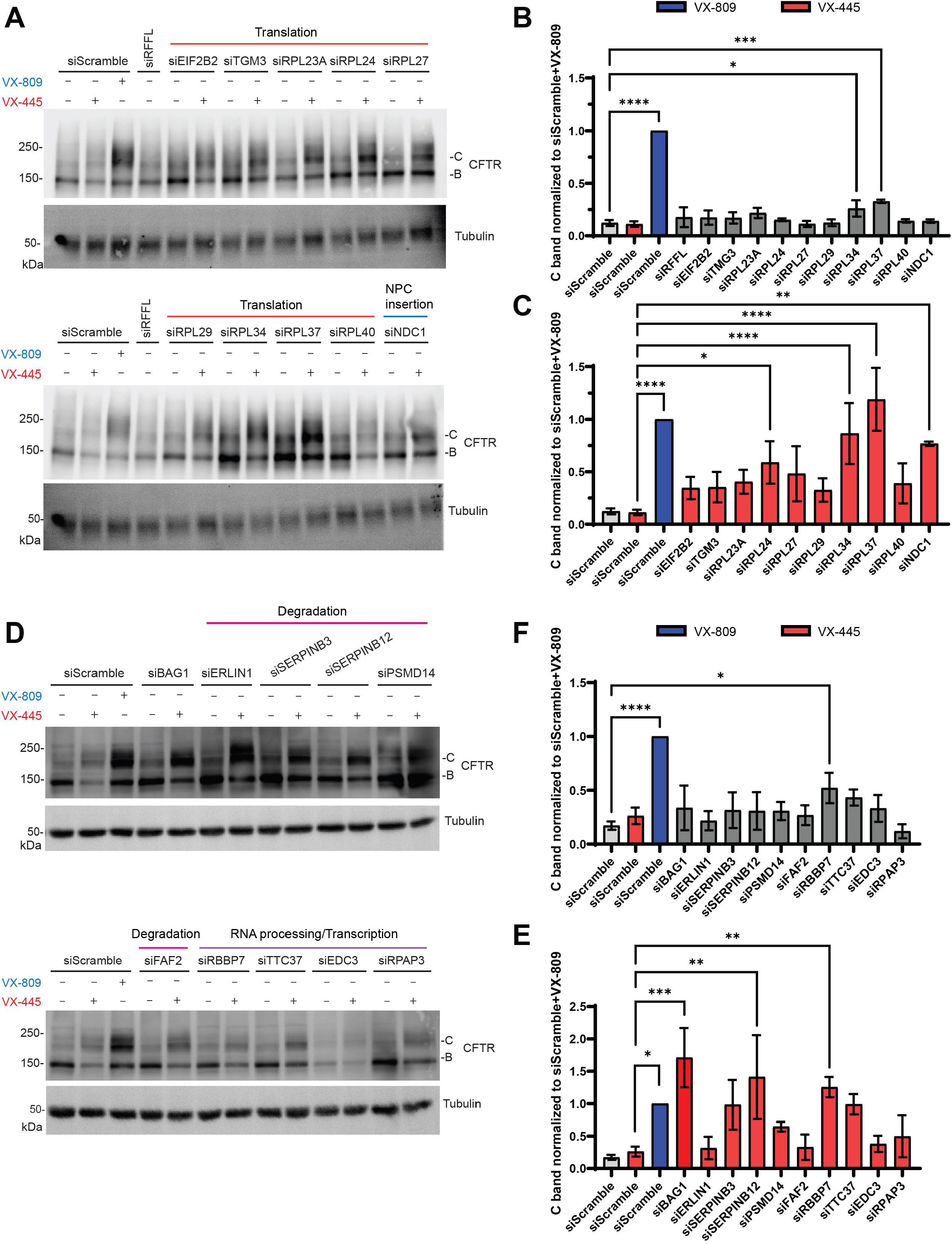
siRNA KD of several RPLs sensitized P67L to VX-445. **A.** Representative Western blot image of P67L showing CFTR expression in Tet-On inducible P67L CFBE cells treated with siRNAs targeting translation factors and nuclear pore complex (NPC) insertion factor at 50 nM for 72 h. DMSO or VX-445 or VX-809 was added at 48 h after siRNA transfection to incubate for 24 h. Scramble siRNA serve as control (n=2-3). **B.** Quantification of band C for P67L treated with siRNAs from A (mean ± SEM). All conditions are normalized to scramble siRNA control treated with VX-809. Mixed effects ANOVA and post-hoc Dunnett multiple comparisons testing was performed to compare each mean to siScramble (p-values: * < 0.05, ** < 0.01, *** < 0.001, and **** < 0.0001) **C.** Quantification of band C for P67L treated with siRNAs and VX-445 from A (mean ± SEM). All conditions are normalized to scramble siRNA control treated with VX-809. Mixed effects ANOVA and post-hoc Dunnett multiple comparisons testing was performed to compare each mean to siScramble + VX-445 (p-values: * < 0.05, ** < 0.01, *** < 0.001, and **** < 0.0001) **D.** Representative Western blot image of P67L showing CFTR expression in Tet-On inducible P67L CFBE cells treated with siRNAs targeting degradation and RNA processing/transcription factors at 50 nM for 72 h. DMSO or VX-445 or VX-809 was added at 48 h after siRNA transfection to incubate for 24 h. (n=2-3). **E.** Quantification of C band of P67L treated with siRNAs from D (mean ± SEM). All conditions are normalized to scramble siRNA control treated with VX-809. Statistical analysis as in B. **F.** Quantification of C band of P67L treated with siRNAs and VX-445 from D (mean ± SEM). All conditions are normalized to scramble siRNA control treated with VX-809. Statistical analysis as in C.

## Discussion

Here we report the first quantitative interactomics of mutant CFTR in response to Elexacaftor/VX-445. Our group has previously compared VX-809 non-responsive and hyper-responsive mutants via AP-MS coupled with TMT labeling to quantitatively explore drug-driven proteostasis network changes^33^. We applied this pipeline to investigate VX-445 and its remodeling of the proteostasis network. We first found the differential rescue of mutant CFTR by VX-445 and compared two rare CFTR mutants, P67L and L206W. Both were hyper-responsive to VX-809 but P67L was only weakly responsive to VX-445 as opposed to L206W. Prior work by Veit and coworkers similarly reported this discrepancy in response to VX-809 and VX-445 for P67L and L206W in CFBE cell-lines^40^. A mild increase in channel function in response to VX-445 in FRT cells stably expressing P67L CFTR as measured by Ussing chamber may also be explained in part by the potentiating activity of VX-445 in addition to a low-level correction^41,42^.

The AP-MS approach showed unique clusters of protein interactors that were impacted differentially by drugs. Some indicated decreased intensities in TMT abundance from DMSO control upon VX-809 or VX-445 treatment to revert the interaction landscape towards WT, while others showed largely unaffected interactions. This validates the capability of our method to capture corrector driven CFTR interactome changes. Our interactomics dataset demonstrated that mutants undergo increased interactions in early CFTR biogenesis pathways responsible for protein quality control such as translation, folding, and degradation in the absence of correctors. VX-809 treatment decreased these interactions indicating proper regulation of translational dynamics and quality control of both P67L and L206W CFTR. VX-445 treatment similarly decreased interactions within the early biogenesis pathways for L206W but did not significantly impact P67L. As the divergent interactome remodeling correlates with the hyper/weak-responsiveness of each mutation, the results reveal potential mutant-specific key checkpoints within pathways such as translation that govern the VX-445 response.

To delineate a causative relationship, we observed several of the protein interactors involved in translation to impact P67L CFTR maturation upon KD via siRNA. KD of large ribosomal subunit proteins such as RPL24, RPL34, and RPL37 drastically improved the overall amount of mature P67L CFTR when simultaneously treated with VX-445. We noted that immature forms of CFTR (B band) were also increased in some KD conditions, especially for siRNA KD of RPL34 and RPL37. Such effects may promote global translation, leading to increased protein in general, including CFTR. While KD of RPL34 has been reported to suppress cell growth and proliferation in cancer cell lines^43,44^, and RPL37 has functions outside the ribosome related to p53 and cell cycle arrest^45,46^, it is unknown what implications their KDs have on global translation dynamics and CFTR. From a purely clinical perspective, patients may benefit from increased absolute levels of mature CFTR on the cell surface. Moreover, trafficking efficiency was increased in these KDs, indicating that increase in C band is not solely due to global translation upregulation but rather VX-445 correction. Additionally, other RPL KDs such as RPL23A and RPL29 showed significant trafficking efficiency improvements, highlighting the role of ribosomal proteins and their involvement in corrector effectiveness. RNAi KD of a selection of RPLs and RPSs was shown to rescue F508del function in CFBE cells^47^, yet none of the RPs from our dataset have been previously assessed to our knowledge. While VX-809 sensitization induced by RPL12 KD has been reported in VX-809 low/non-responders F508del, G85E, and A455E^48^, our study reports the first set of siRNA KDs that induce VX-445 sensitization of a low/non-responder.

Based on the KD findings, we propose that translational dynamics are dysregulated in P67L, which could result in insufficient conformational sampling of mutant CFTR for VX-445 binding. The binding site of VX-445 identified in the cryo-EM CFTR structure indicates the N-terminal lasso motif participates in VX-445 attachment^19^. The lasso motif in CFTR may be impaired by P67L^37^, which may explain the loss in effectiveness of VX-445 for P67L. CFTR folds co-translationally^49,50^ in a highly fine-tuned manner where mutations or premature folding of subdomains can disrupt conformations of the overall domain^51,52^. Interestingly, KD of eIF3a, a subunit of the eIF3 initiation factor, rescued CFTR folding in various mutants by slowing translation^53^. Blunting translational rate by altering ribosomal velocity via RPL12 KD has been shown to rescue F508del maturation^48^. Lastly, early co-translational misfolding of P67L may result in ribosome stalling and trigger ribosomal quality control to terminate translation^54^. Based on these previously reported findings, together with observations presented here, we suspect that alterations in ribosomal translational dynamics resulting from KD of specific RPLs (potentially slowed translation) may allow sufficient time for P67L domains to fold into metastable states that facilitate productive binding of VX-445 (**Figure 5**). Once VX-445 is properly bound to P67L, remaining subdomains may translate and fold over the existing bound and stable form of premature CFTR, allowing evasion of subsequent quality control checkpoints. In contrast, F508del or L206W likely do not require translational tampering for VX-445 to bind. The structural defects introduced by these mutations are distant from the cryo-EM binding site and mutation induced co-translational misfolding may occur later than for P67L.

**Figure 5.**
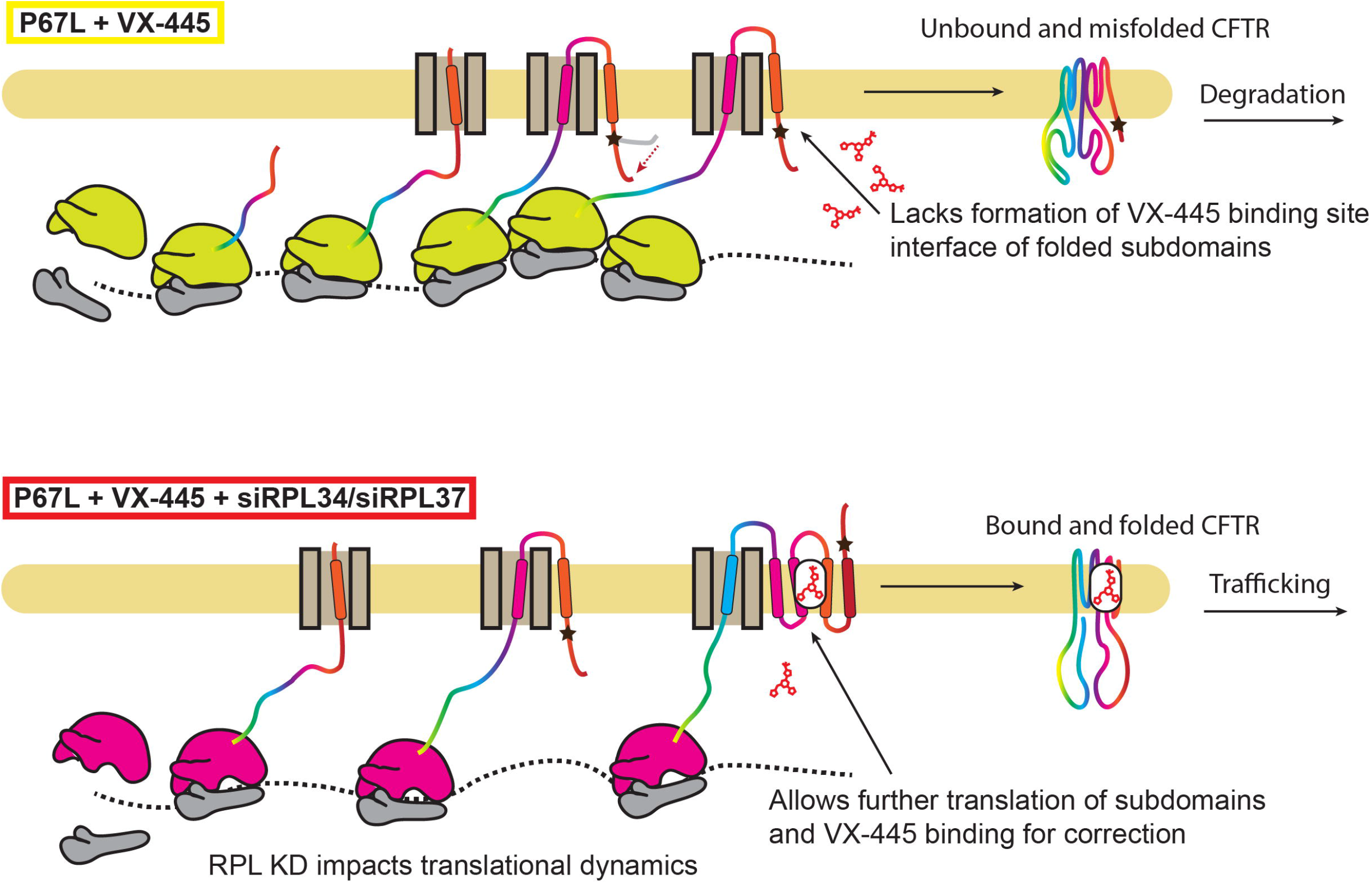
Co-translational misfolding induced by P67L may cause ribosome stalling early during translation. Alterations in ribosomal translational dynamics resulting from RPL KD, may allow further translation and allow subdomains to fold for proper binding of VX-445.

Another possible explanation for the VX-445 sensitization induced by RPL KD may involve individual functions of RPLs outside the ribosome. Such functions include tumorigenesis, immune signaling, tissue development and roles in multiple diseases such as autism and Parkinson’s disease^55^. Therefore, ribosomal proteins may serve functions other than ribosomal binding that involves CFTR protein quality control. Our siRNA screen highlights ways in which structural proteins located at diverse positions on the ribosome^56^ may nonetheless produce net similar effects on P67L CFTR biogenesis. Alternatively, disrupting the relative proportion of ribosomal subunits may result in a smaller subpopulation of functioning ribosomes and a larger subpopulation of altered ribosomes that exhibit aberrant behavior regardless of the exact identity of the missing RPL. Further experiments to assess translation speed, polysome composition, and ribosomal footprinting in response to RPL KDs will be required to distinguish these possibilities.

Most hits identified from degradation/folding siRNA pathway did not induce strong sensitization of P67L to VX-445, although it is well-established in the literature that inhibition or KD of pro-degradation factors can rescue F508del CFTR ^30,32,57–59^. In semblance to possible mutation specific VX-445 binding site disruption, different class II mutants may depend on altered protein quality control and subsequent degradation pathways as reported previously^33^ and require mutation specific bypassing of quality control checkpoints^60^. Nonetheless, SERPINB12 and RBBP7 sensitized P67L to VX-445. SERPINs are trypsin-like serine-proteases inhibitors and SERPINB12 may regulate transcription via RNA polymerase II binding^61,62^. RBBP7 is part of chromatin remodeling complexes and is implicated in transcriptional repression via histone deacetylation^63^. Histone deacetylases (HDACs) have been shown to improve F508del expression upon inhibition with HDAC inhibitors^64^. Thus, the sensitization displayed by SERPINB12 and RBBP7 may be explained by the epigenetic influence on CFTR translation dynamics^65^.

We included RFFL and BAG1 in our panel as they were previously reported to rescue P67L and F508del CFTR upon KD, respectively^66,67^. While we did not observe significant rescue of P67L with RFFL or BAG1 KD, BAG1 KD greatly sensitized P67L to VX-445. BAG1 was reported to promote proteasomal degradation of misfolded proteins in a ubiquitination dependent manner^68^. However, BAG1 was also reported to confer stability to F508del CFTR and thus KD leads to destabilization and subsequent loss of F508del CFTR^69^. Despite the conflicting literature, we suspect that depletion of BAG1 results in a larger amount of residual misfolded P67L CFTR that is available for correction by VX-445.

Our interactomics profiling provides a better understanding of the VX-445 driven protein-interactions involved in CFTR biogenesis and may indicate potential targets for further therapeutic investigation. While we focused on translational factors in this study, a rich dataset is available for mining of other potential hits that may further fuel discovery. Our approach can be applied to uncover dysregulated proteostasis states in numerous protein misfolding diseases that are linked to proteostasis deformations^70^. Furthermore, the mechanistic insight regarding corrector function within cellular processes can aid in optimizing combination treatments for patients to augment corrector effectiveness as shown here for P67L. Similar approaches can be taken for Elexacaftor or Trikafta refractory mutants to benefit the currently unserved ~6% of CF patients ineligible for Trikafta.

## Acknowledgments

We thank Dr. Guido Veit and Dr. Gergely Lukacs (McGill University) for sharing CFBE41o-TetON CFTR expressing cells. We thank Dr. Katherine Oliver (Emory University) for helpful discussion and members of the Plate lab for their critical reading and feedback on this manuscript.

## Author Contributions

Conceptualization (L.P., C.M.P.S., E.F.M., M.K.); Investigation (M.K., E.F.M, B.T., D.J.); Data curation & Formal Analysis (M.K., E.F.M, B.T., D.J., J. H., A.R., L.P); Funding acquisition (L.P., E.F.M, C.M.P.S., E.J.S.); Methodology (L.P., J.S.H, A.R, E.J.S); Project administration (L.P., E.J.S.); Writing – original draft (M.K, L.P.); Writing – review & editing (all authors)

## Funding and additional Information

This work was funded by T32 GM065086 (NIGMS) and F31 HL162483 (NHLBI) (EFM); the Cystic Fibrosis Foundation Postdoctoral Fellowship (SABUSA19F0) (CMPS); R01 HL139876 (NHLBI) (EJS); R35 GM133552 (NIGMS) (LP); and Vanderbilt University funds. The content is solely the responsibility of the authors and does not necessarily represent the official views of the National Institutes of Health.

## Data Availability

The mass spectrometry proteomics data have been deposited to the ProteomeXchange Consortium via the PRIDE^71^ partner repository with the dataset identifier PXD039773.

## Supporting Information

This article contains supporting information^31,72^.

## Methods

### Plasmids and antibodies

Plasmids used for transient transfection expressed WT or mutant CFTR in the pcDNA5/FRT vector. Anti-CFTR antibodies used for detection were 217 and 596 (provided by J. Riordan, University of North Carolina, Chapel Hill, North Carolina; http://cftrantibodies.web.unc.edu/) each at 1:1000 and 1:500 working dilutions in immunoblotting buffer (5% bovine serum albumin [BSA] in Tris-buffered saline, pH 7.5, 0.1% Tween-20, and 0.1% NaN_3_) respectively. Antibodies used for immunoprecipitation was mAB 24-1 (ATCC HB-11947) purified from B lymphocyte hybridoma cells using a recombinant Protein G–Sepharose 4B Affinity Column on an ÄKTA start protein purification system (GE Life Science Product # 29022094). Secondary antibodies used were goat anti-mouse StarbrightB700 (Bio-Rad), anti-mouse IgG HRP conjugate (Promega) anti-rabbit rhodamine–conjugated tubulin (Bio-Rad).

### Cell culture

Human embryonic kidney 293T (HEK293T) cells were cultured in Dulbecco’s modification of Eagle’s medium (DMEM, Corning) supplemented with 10% fetal bovine serum (FBS, Gibco), 1% L-glutamine (200 mM, Gibco), and 1% penicillin/streptomycin (10,000 U; 10,000 μg/mL, Gibco).

Tet-On inducible human bronchial epithelial cells were generously provided by Dr. Guido Veit and Dr. Gergely Lukacs, McGill University^39^. F508del-CFTR (iF508del-CFBE), P67L-CFTR (iP67L-CFBE), WT-CFTR (WT-CFBE) and parental CFTR null (CFBE41o^-^) were cultured in minimum essential medium (MEM, Gibco) supplemented with 10% FBS (Gibco), 1% HEPES (1 M, Gibco), 1% L-glutamine (200 mM, Gibco). For selection of inducible cells, media was further supplemented with 200 μg/mL G418 (Invivogen) and 3 μg/mL puromycin (Invivogen). For immunoblotting experiments, CFBE cells were allowed to differentiate for at least 72 h after full confluency. Induction of CFTR expression was performed with 500ng/L doxycycline (Fisher Bioreagents) for at least 72 h before collection.

Fischer rat thyroid (FRT) cell lines expressing P67L or L206W CFTR, established by Sorscher laboratory^35^, were cultured in Coon’s modification of Nutrient Mixture F-12 Ham (Sigma F6636) complemented with 2.68 g/l sodium-bicarbonate in the presence of 5% fetal bovine serum (Gibco). The media was supplemented with 100 μg/ml Hygromycin B (Invitrogen, 10687010) for expansion cell culture to serve as selection pressure. The cells were maintained in 37C° humidifier incubator at 5% CO2 – 95% air.

### Immunoblotting

HEK293T cells were transiently transfected with CFTR plasmids via calcium-phosphate transfection^73^. Cells were exposed to respective drug treatments at 3 μM for 24 h before collection. Cells were rinsed with phosphate-buffered saline (PBS) and lysed on plate with 600 μL of TNI buffer (50 mM Tris-base, 150 mM NaCl, pH 7.5, and 0.5% IGEPAL CA-630, EDTA-free protease inhibitor cocktail [Roche]) rocking at 4°C for 20 min. Cells were harvested by scraping and sonicated for 3 min and centrifuged at 18,000×g for 30 min. Resulting lysate was normalized with Bio-Rad protein assay dye. Samples were denatured in 2.4× Laemmli buffer with 10 mM dithiothreitol, resolved by 8% SDS–PAGE, and transferred to PVDF membranes (Millipore). Membranes were blocked in 5% milk in Tris-buffered saline, pH 7.5, 0.1% Tween-20 (TBS-T) at RT for 30 min. Membranes were washed with TBS-T and probed in primary antibodies overnight at 4°C. After three washes with TBS-T, membranes were probed with secondary antibody at RT for 30 min. Membranes were washed three times with TBS-T and imaged using a ChemiDoc MP Imaging System (Bio-Rad). HRP conjugate blots were developed with chemiluminescent substrate (Clarity Western ECL, Bio-Rad). Quantification was performed using ImageLab (Bio-Rad). Immunoblot quantification data statistical differences were computed with a two-tailed paired t test (p-values: * < 0.05, ** < 0.01, *** < 0.001, and **** < 0.0001).

### Ussing chamber measurements

Fischer Rat Thyroid (FRT) cells carrying P67L or L206W mutation were seeded and cultured on Transwell permeable supports (Corning). The cells formed polarized monolayers with a minimum Rt of 400 Ω× cm^2^ in 5 days under liquid/liquid condition. Monolayers were treated with VX-809, VX-661 and/or VX-445 (SelleckChem) for 48 hours from apical and basolateral side prior Ussing assay. On the day of assay, monolayers were mounted onto EasyMount Ussing Chamber System (Physiologic Instruments) and bathed apically in low chloride Ringer’s solution: 140 mM Na-gluconate, 1.2 mM NaCl, 25 mM NaHCO_3_, 3.33 mM KH_2_PO_4_, 0.83 mM K_2_HPO_4_, 1.2 mM CaCl_2_, 1.2 mM MgCl_2_, and 10 mM D-glucose (pH 7.4) whereas basolaterally the Na-gluconate was replaced by 120 mM NaCl to maintain chloride gradient. The bathing solutions were maintained at 37 °C and stirred by bubbling through 5% CO_2_/95% O_2_. Once current stabilized, 100 μM amiloride (MilliporeSigma) was applied to both apical and basolateral sides to inhibit epithelial sodium channel ENaC. CFTR protein was activated by applying 5 μM forskolin (MilliporeSigma) to both sides followed by VX-770 (SelleckChem) apical addition at 5 μM concentration to investigate gating abnormalities of CFTR variants. At the end of each experiment, CFTR (inh)-172 (10 μM; MilliporeSigma) was applied to the apical side to inhibit CFTR-mediated current. Short circuit current (Isc) was measured under voltage clamp condition and the changes induced by activator or inhibitor were calculated and expressed as mean ± standard deviation.

### Co-Immunoprecipitation and multiplexed LC-MS/MS

Co-immunoprecipitation (co-IP) of CFTR bound with interactors was carried out as described previously^74^. Briefly, cell lysates were pre-cleared with 4B Sepharose beads (Sigma) at 4°C for 1 h while rocking. Precleared lysates were then immunoprecipitated with Protein G beads complexed to 24-1 antibody (6 mg antibody/mL of beads) overnight at 4°C while rocking. Beads were washed three times with TNI buffer, twice with TN buffer, and frozen at –80°C for at least 1 h. Proteins were then eluted with shaking at 37°C for 1 h with elution buffer (0.2 M glycine, 0.5% IGEPAL CA-630, pH 2.3). Elution steps were repeated once and combined to immediately neutralize with ammonium bicarbonate solution.

MS sample preparation of co-IP samples were performed as described previously^33^. Briefly, samples were precipitated in methanol/chloroform. Precipitated pellet was rinsed and dried to reconstitute in 1% Rapigest SF (Waters). Samples were reduced with 5 mM TCEP (Sigma), alkylated with 10 mM iodoacetamide (Sigma), and digested with 0.5 μg of trypsin (Sequencing Grade, Promega, or Pierce) overnight in 50mM HEPES (pH 8.0) at 37°C with shaking. Digested peptides were labeled with TMT 11-plex reagents (Thermo Fisher). TMT-labeled samples were pooled and acidified with MS-grade formic acid (Sigma) to remove cleaved Rapigest SF via centrifugation. Samples were concentrated using a SpeedVac (Thermo Fisher) and resuspended in buffer A (95% water, 4.9% acetonitrile, and 0.1% formic acid). Sample was then loaded onto a triphasic MudPIT^75^ column using a high-pressure chamber.

LC-MS/MS analysis was performed on an Exploris 480 (Thermo Fisher) mass spectrometer equipped with an UltiMate3000 RSLCnano System (Thermo Fisher) as described previously^33^. Briefly, 10 μl sequential injections of 0, 10, 30, 60, and 100% buffer C (500 mM ammonium acetate in buffer A) were performed followed by a final injection of 90% buffer C with 10% buffer B (99.9% acetonitrile, 0.1% formic acid). Each step consisted of a 90-min gradient from 4 to 40% B with a flow rate of either 300 or 500 nL/min, followed by a 15-min gradient from 40 to 80% B with a flow rate of 500 nL/min on a 20-cm fused silica microcapillary column (ID 100 μM) ending with a laser-pulled tip filled with Aqua C18, 3 μm, 100 Å resin (Phenomenex). Electrospray ionization was performed directly from the analytical column by applying a voltage of 2.0 or 2.2 kV with an inlet capillary temperature of 275°C. Data-dependent acquisition of MS/MS spectra was performed by scanning from 300 to 1800 m/z with a resolution of 60,000 to 120,000. Peptides with an intensity above 1.0E4 with charge state 2–6 from each full scan were fragmented by HCD using normalized collision energy of 35 to 38 with a 0.4 m/z isolation window, 120 ms maximum injection time at a resolution of 45,000, scanned from 100 to 1800 m/z or defined a first mass at 110 m/z and dynamic exclusion set to 45 or 60s and a mass tolerance of 10 ppm.

Peptide identification and TMT-based protein quantification was carried out using Proteome Discoverer 2.4 as described previously^33^. MS/MS spectra were extracted from Thermo XCaliber .raw file format and searched using SEQUEST against a UniProt human proteome database (released 03/25/2014) containing 20,337 protein entries. The database was curated to remove redundant protein and splice-isoforms and supplemented with common biological MS contaminants. Searches were carried out using a decoy database of reversed peptide sequences and the following parameters: 10 ppm peptide precursor tolerance, 0.02 Da fragment mass tolerance, minimum peptide length of six amino acids, trypsin cleavage with a maximum of two missed cleavages, static cysteine modification of 57.0215 Da (carbamidomethylation), and static N-terminal and lysine modifications of 229.162932 Da (TMT 11-plex). SEQUEST search results were filtered using Percolator to minimize the peptide false discovery rate to 1% and a minimum of two peptides per protein identification. TMT reporter ion intensities were quantified using the Reporter Ion Quantification processing node in Proteome Discoverer 2.4 and summed for peptides belonging to the same protein.

### Interactor filtering and statistical analysis

A total of 6 TMT-11plex sets of samples were analyzed via LC-MS/MS over six separate mass spectrometry runs. Interactors co-immunoprecipitated with CFTR were filtered by comparing against a mock transfection control (tdTomato). The log_2_ fold change of total peptide normalized abundances of each protein over mock were computed within each run. The 6 separate log_2_ fold change representing individual MS runs were then averaged to yield the consensus log_2_ fold change over mock values for each protein and a paired two-tailed t test was used to calculate the p-value (**Table S1 and S2**).

To filter for true interactors of CFTR, a curved filter combining log_2_ fold change and p-value was used as described previously^76^. Briefly, the histogram of log_2_ fold changes over mock were fitted to a Gaussian curve using a nonlinear least-square fit to determine the standard deviation σ. Fold change cutoff for interactors was set to 1 σ. A reciprocal curve with the equation y > c/(x - x_0_), where y = p-value, x = log_2_ fold change, x_0_ = fold change cutoff (1 σ), and c = the curvature (c = 0.8) was used to filter interactors in each condition. These interactors were pooled to generate a master list of true CFTR interactors. The abundance values of these interactors in each condition were then normalized to WT CFTR. Briefly, each abundance value of an individual run was log_2_ transformed and averaged to yield the consensus log_2_ grouped abundance. These values for each condition were subtracted by WT log_2_ grouped abundance for each protein and subsequently subtracted by WT CFTR log_2_ grouped abundance to yield the log_2_ fold change of protein abundances against the WT standardized to CFTR levels across conditions.

For aggregate pathways statistics in violin plots, we used a one-way ANOVA with Geisser-Greenhouse correction and post-hoc Tukey multiple comparisons testing to evaluate the statistically significant difference between conditions for a given pathway.

### Hierarchical clustering and pathway assignment

Hierarchical clustering of proteins in the master list of true CFTR interactors was performed with a custom R script. Briefly, the WT CFTR normalized abundances were converted to an Euclidean distance matrix and clustered using Ward’s minimum variance method.

The proteins in our master list were searched against a previously reported dataset^31^. This step classified all previously identified interactors. Novel interactors were assigned with pathways by searching the UniProt database. Common MS contaminants were removed by querying list of proteins against CRAPome 2.0^72^.

### siRNA knockdown

CFBE cells were transfected with 50nM siRNA at 60% confluence using Lipofectamine RNAiMax transfection reagent, following the manufacturer’s protocols (Invitrogen). Cells were also induced with 500ng/L doxycycline (Fisher bioreagents) at the time of transfection and collected after 72 h.

### RT-qPCR

RNA from siRNA knockdown samples were isolated with Quick-RNA Miniprep kit (Zymo). cDNA library was generated with 500 ng of RNA, 250 ng random hex primers (Integrated DNA Technologies), 250 ng poly-T primers (Integrated DNA Technologies), and M-MLV reverse transcriptase (Promega). Quantitative PCR was performed with iTaq Universal SYBR green supermix (Bio-Rad) on the CFX96 Touch Real-Time PCR Detection System (Bio-Rad). Measurements were taken in 3 technical replicates and normalized to GAPDH control.

