## Supporting Information for "Elexacaftor/VX-445-mediated CFTR interactome remodeling reveals differential correction driven by mutation-specific translational dynamics"

for

| | |
|---|---|
| Supporting Figures S1-S7 | p S2-S8 |
| Supporting Tables S1-S3 | p S9 |
| References | p S10 |

**Supporting Figures**

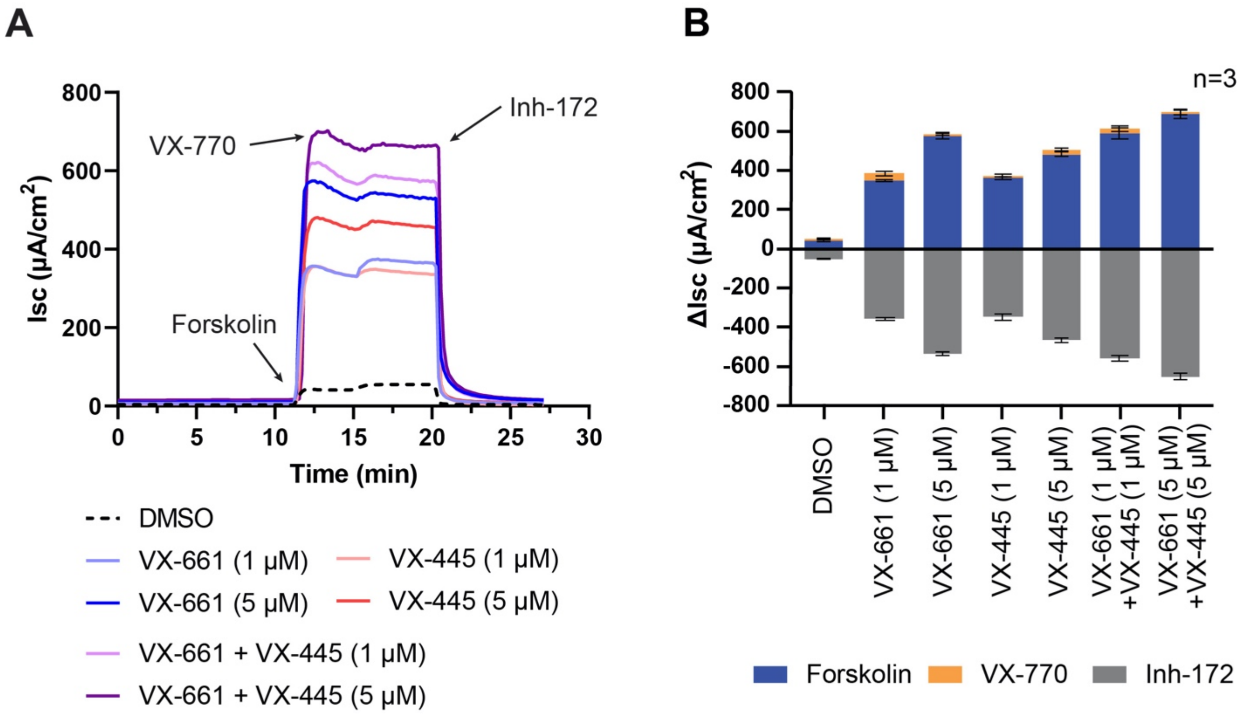


**Figure S1. FRT cells stably expressing L206W CFTR treated with VX-661 or VX-445 shows concentration dependent functional rescue.**

**A.** Ussing chamber measurements on FRT cells stably expressing L206W. Data showing short-circuit current (Isc) after luminal stimulation with forskolin (5 µM) and VX-770 (5 µM) and inhibition by CFTR inh-172 (10 µM).

**B.** Bar graph representation of the acute treatment induced Isc changes are expressed as mean±SD across 3-4 biological replicates.

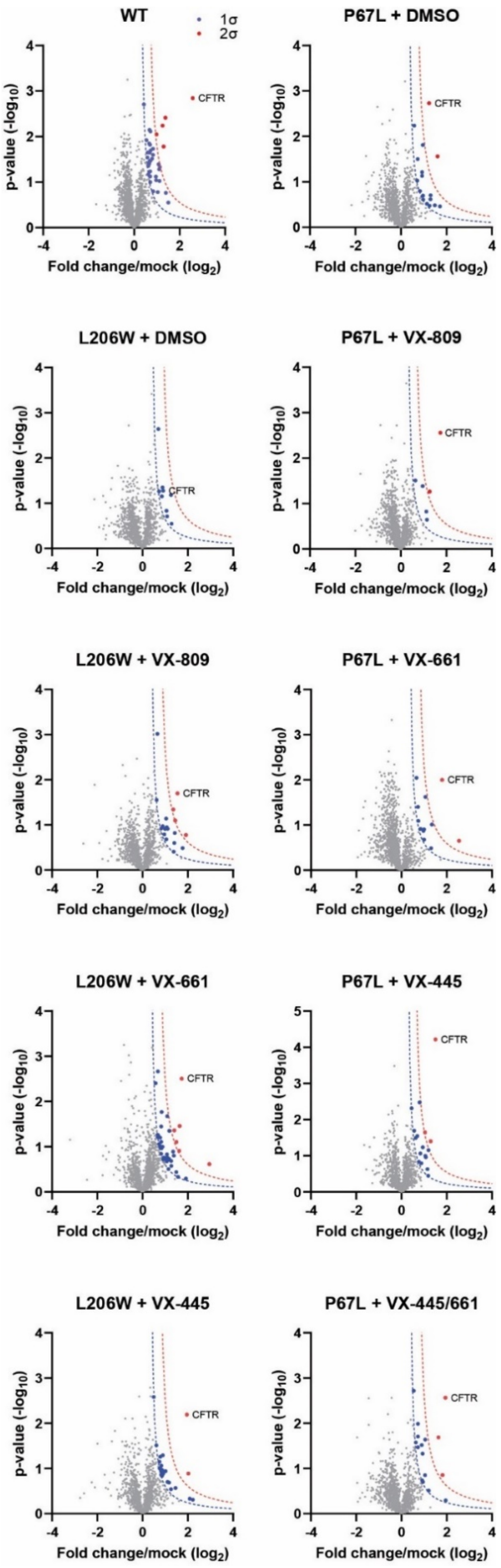


**Figure S2.** Filtering for true interactors. Volcano plots of each condition is shown where the $\log_2$ fold changes of TMT intensities over mock transfection control are shown on the x-axis and $-\log_{10}$ p-values are shown on the y-axis. Confidence interactors are filtered for one standard deviation away from the normal distribution of all. These confidence interactors from each condition were pooled into a master list of interactors for further analysis (**Table S2)**.

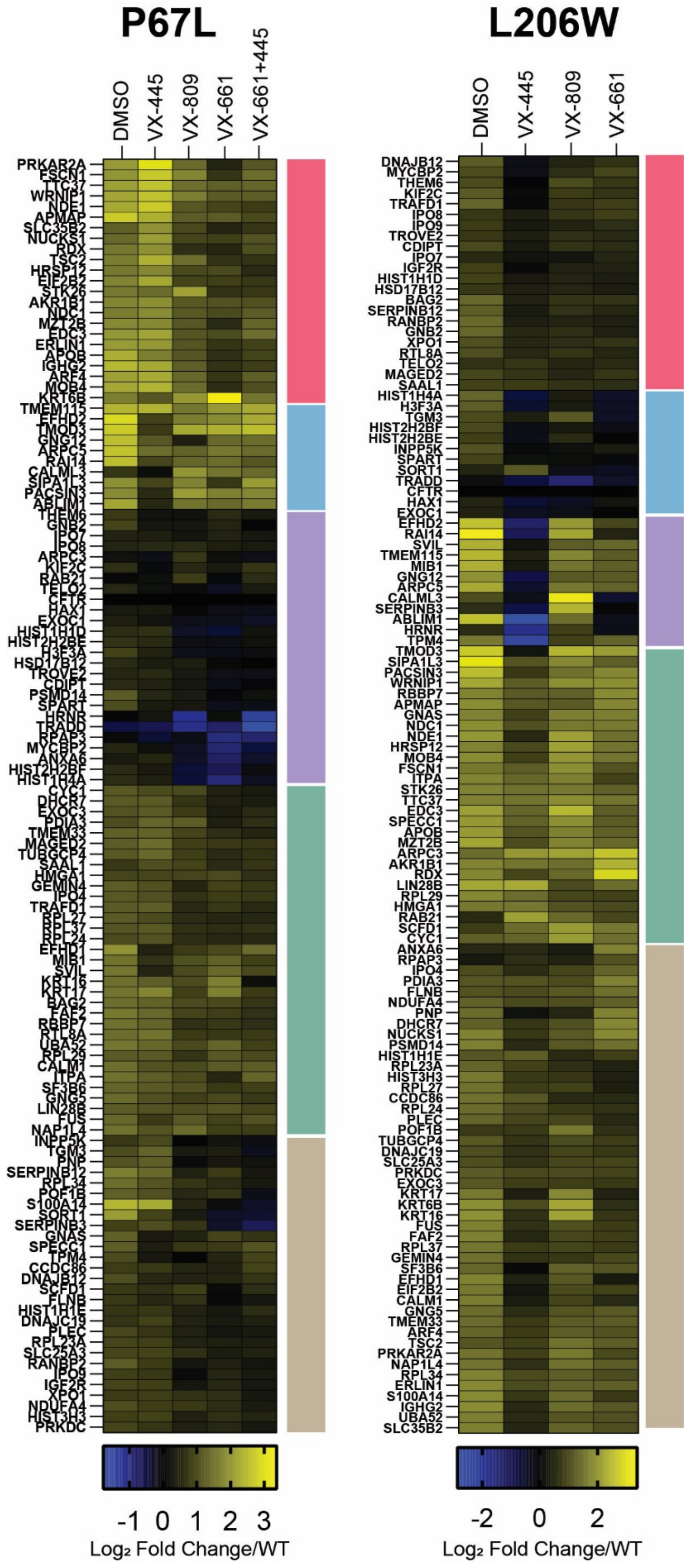


**Figure S3.** Hierarchical clustering of interactors grouped based on intensity similarity where each bar is an interacting protein and yellow indicates increased interaction with CFTR when compared to WT. All conditions in the 11-plex TMT experiment are shown for each mutant. Each hierarchical cluster is represented as colored blocks.

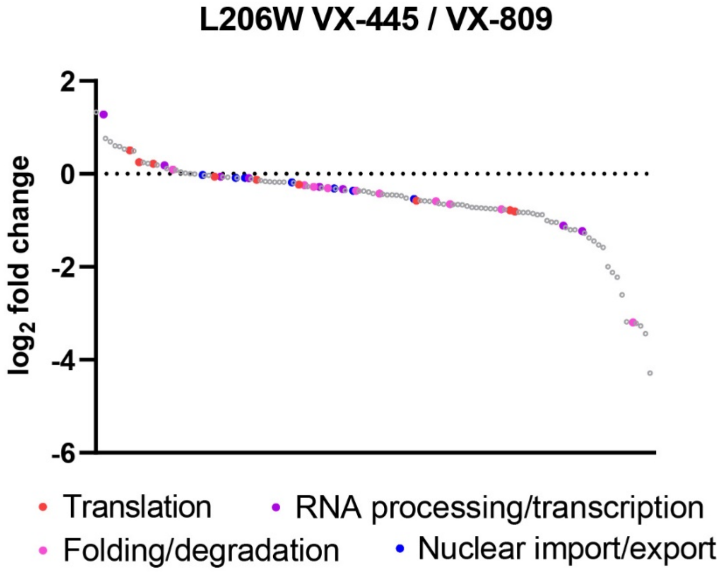


**Figure S4.** Waterfall plot directly comparing VX-445 to VX-809 treatment for L206W. Higher $log_2$ fold change indicates higher interaction in VX-445 treated L206W than that of VX-809 treated L206W. Distribution of hits determined from P67L are more arbitrarily distributed as noted by the red and pink data points.

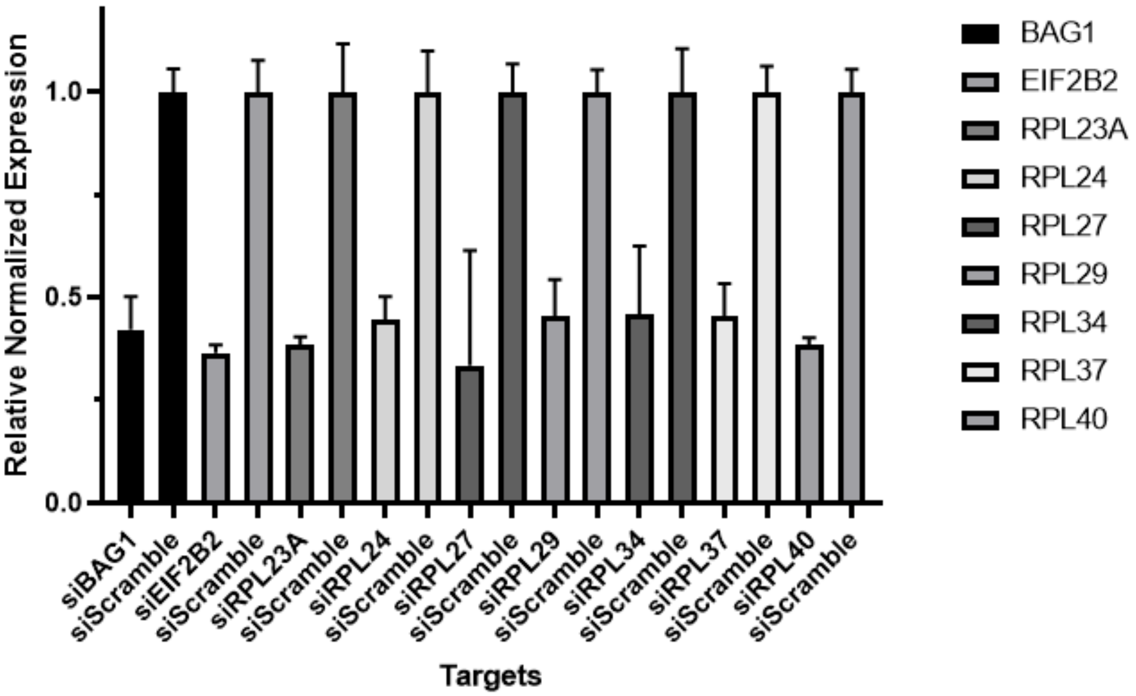


**Figure S5.** Expression of siRNA KD targets in inducible P67L CFBE cells normalized to siScramble control as measured via RT-qPCR. Expression was normalized to GAPDH within each sample prior to siScramble normalization.

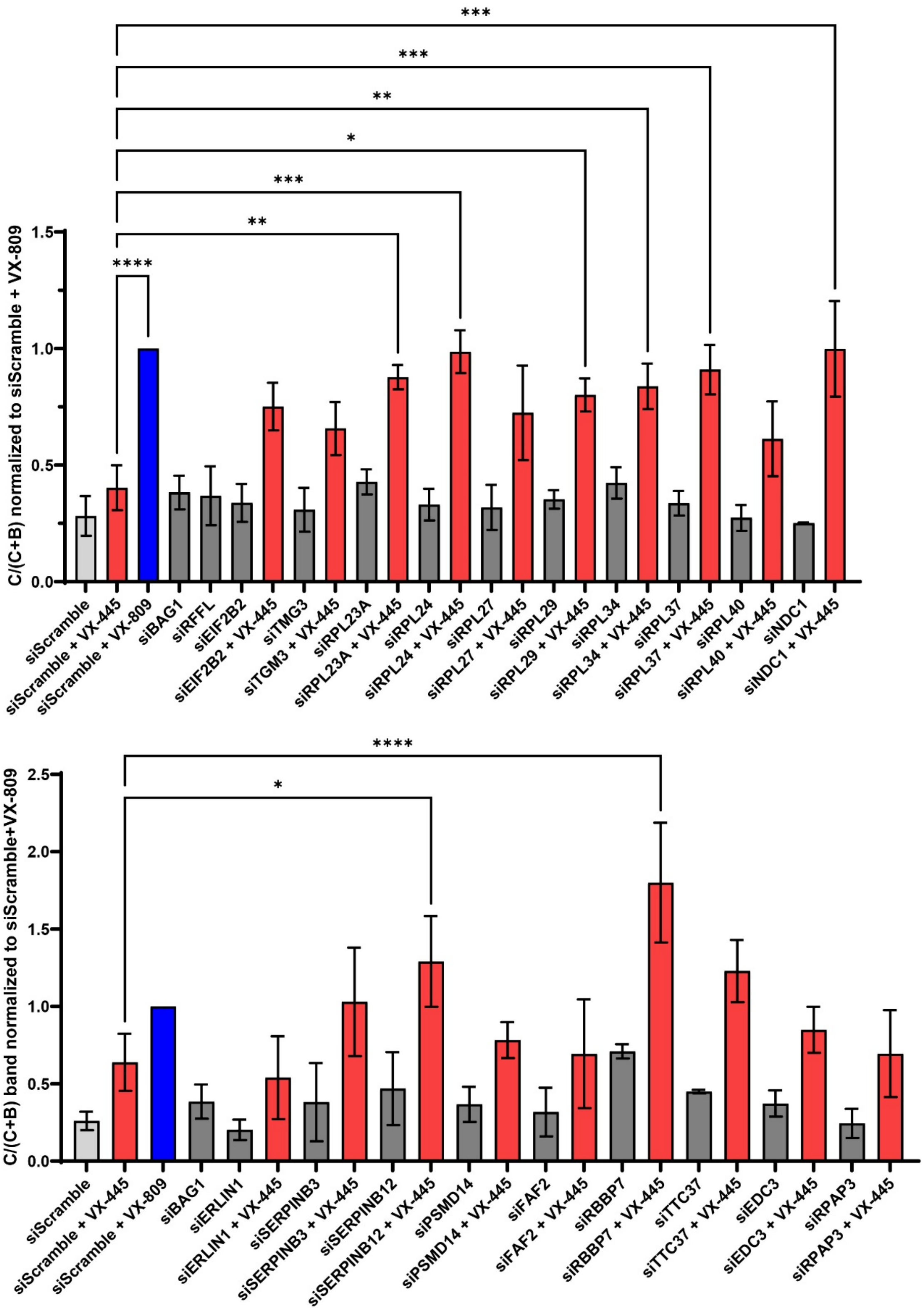


**Figure S6.** Quantification of trafficking efficiency of CFTR from western blot images of Tet-On inducible P67L CFBE cells treated with siRNAs ± VX-445 (mean ± SEM). All conditions are normalized to scramble siRNA control treated with VX-809. Statistical differences were computed via Mixed effects ANOVA and post-hoc Dunnett multiple comparisons testing was performed to compare each mean to siScramble + VX-445 (p-values: * $< 0.05$, ** $< 0.01$, *** $< 0.001$, and **** $< 0.0001$).

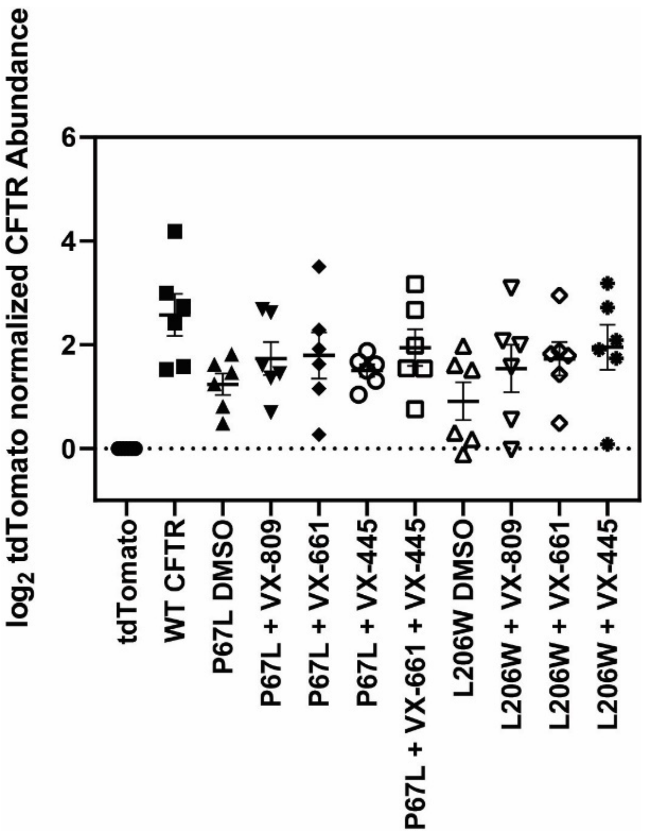


**Supplemental Figure S7.** $Log_2$ value of CFTR abundances in each TMT 11-plex channel normalized to tdTomato (mock transfection). Each datapoint represents a biological replicate. (n=6)

**Supporting Tables**

**Table S1.** Outline of experimental conditions labeled with TMT 11-plex reagents. Six individual mass spectrometry experiments were performed.

**Table S2.** List of filtered interactors displaying a $log_2$ fold enrichment (CFTR/control) greater than one standard deviation based on the distribution of identified proteins. Each CFTR variant and corrector treatment was compared to the control within each replicate to average across 6 replicates and normalize to WT enrichment. Proteins were assigned to corresponding pathways by comparing with previously published dataset[1] and searching the UniProt database. Common MS contaminants were removed by querying list of proteins against CRAPome 2.0[2].

**Table S3.** List of prioritized siRNA targets as determined from waterfall plot (**Figure 3D**).
